# Increased Colonic Expression of ACE2 Associates with Poor Prognosis in Crohn’s disease

**DOI:** 10.1101/2020.11.24.396382

**Authors:** Takahiko Toyonaga, Kenza C. Araba, Meaghan M. Kennedy, Benjamin P. Keith, Elisabeth A. Wolber, Caroline Beasley, Erin C. Steinbach, Matthew R. Schaner, Animesh Jain, Millie D. Long, Edward L. Barnes, Hans H. Herfarth, Kim L. Isaacs, Jonathan J. Hansen, Muneera Kapadia, José Gaston Guillem, Mark J. Koruda, Reza Rahbar, Tim Sadiq, Ajay S. Gulati, Praveen Sethupathy, Terrence S. Furey, Camille Ehre, Shehzad Z. Sheikh

## Abstract

**Background and Aims:** The host receptor for SARS-CoV-2, angiotensin-converting enzyme 2 (ACE2), is highly expressed in small intestine. Our aim was to study colonic ACE2 expression in Crohn’s disease (CD) and non-inflammatory bowel disease (non-IBD) controls. We hypothesized that the colonic expression levels of ACE2 impacts CD course.

**Methods:** We examined the expression of colon *ACE2* using RNA-seq and quantitative (q) RT-PCR from 69 adult CD and 14 NIBD control patients. In a subset of this cohort we validated ACE2 protein expression and localization in formalin-fixed, paraffin-embedded matched colon and ileal tissues using immunohistochemistry. The impact of increased *ACE2* expression in CD for the risk of surgery was evaluated by a multivariate regression analysis and a Kaplan-Meier estimator. To provide critical support for the generality of our findings, we analyzed previously published RNA-seq data from two large independent cohorts of CD patients.

**Results:** Colonic *ACE2* expression was significantly higher in a subset of adult CD patients (ACE2-high CD). IHC in a sampling of ACE2-high CD patients confirmed high ACE2 protein expression in the colon and ileum compared to ACE2-low CD and NIBD patients. Notably, we found that ACE2-high CD patients are significantly more likely to undergo surgery within 5 years of diagnosis, with a Cox regression analysis finding that high *ACE2* levels is an independent risk factor (OR 2.18; 95%CI, 1.05-4.55; p=0.037).

**Conclusion:** Increased intestinal expression of ACE2 is associated with deteriorated clinical outcomes in CD patients. These data point to the need for molecular stratification that may impact CD disease-related outcomes.

## Introduction

Crohn’s disease (CD) is a chronic inflammatory condition of the intestinal tract affecting millions of people worldwide ^1–3^. CD patients frequently require immunosuppressant medications, which can increase the risk of infection, including for respiratory diseases such as influenza and pneumonia. COVID-19 infections are increasing world-wide (https://coronavirus.jhu.edu). A significant number of patients present with gastrointestinal symptoms and high levels of viral RNA in the stool have been detected. This has led the IBD research community to investigate molecules associated with SARS-CoV2 infectivity with an emphasis on its cognate receptor ACE2. ACE2 is essential for viral entry into epithelial cells and is abundantly expressed in the lung and intestinal epithelium, with markedly higher expression in the small intestine under normal conditions. Expression of two mucosa-specific serine proteases, TMPRSS2 and TMPRSS4, also promote SARS-CoV-2 virus entry into host cells. In the small intestine, levels of expression of *ACE2* in patients with CD are dependent on inflammation status and the specific anatomical location^4^.

Disease presentation and progression within CD is highly heterogeneous in location, severity of inflammation, and other phenotypes. Current CD clinical classifications fail to accurately predict disease-related outcomes. Defining on a molecular basis subsets of IBD patients with similar severity is essential for developing guidelines for the use of standard IBD therapies. Recently, Suárez-Fariñas et al. showed high small bowel enterocyte brush border expression of ACE2 and TMPRSS2. IBD medications, both biologic and non-biologic, did not significantly impact the expression of both genes in the uninflamed small intestine^5^. However, Potdar et al. revealed that within CD, small bowel ACE2 was reduced in patients subsequently developing complicated disease and that its expression was restored in responders to biologic therapy^6^. However, there remain three major gaps in our knowledge not addressed by recent studies reporting on ACE2 expression and its association with clinical IBD. First, the role of colonic ACE2 in predicting disease course in CD remains unstudied. Second, how the expression of ACE2 relates to that of other genes, as determined by unbiased transcriptomics, needs to be elucidated. Finally, the relationship of colonic and ileal ACE2 expression in the same patient and its association to disease outcome is unknown.

In this current study, we show that expression of *ACE2*, as well as *TMPRSS2* and *TMPRSS4*, are highly variable in the intestines of adult and pediatric patients with CD, and that their expression levels associate longitudinally with IBD outcome. Our work reveals a novel connection between colonic ACE2 expression and CD-associated clinical outcomes. These findings motivate future studies that focus on differences in ACE2 regulation between ileum and colon in Crohn’s disease and also on whether colonic epithelial SARS-CoV-2 infectivity is greater in the ACE2-high subtype of patients.

## Materials and Methods

### Subjects, Samples, and Clinical Information

Colonic mucosa was obtained from surgically resected colon specimens from patients with an established diagnosis of CD between February 2012 and Jan 2018. All samples were collected from disease-unaffected regions without macroscopic inflammation and were from ascending colon. Clinical information was collected from medical records up to 5 years after CD diagnosis.

Two Independent cohorts of adult CD and treatment-naïve pediatric CD samples were downloaded from GEO (accession numbers GSE57945 and GSE137344). Pediatric CD samples from GSE57945 were processed as described previously ^12^.

### RNA isolation, sequencing, and processing

Adult samples from UNC hospitals were isolated and sequenced as previously described ^8^. Briefly, RNA was isolated using the Qiagen RNeasy Mini Kit following the manufacturer’s protocol, and RNA purity was assessed with Thermo Scientific NanoDrop 2000. RNA-seq libraries were prepared using the Illumina TruSeq polyA+ Sample Prep Kit. Paired-end (50 bp) sequencing was performed on the Illumina HiSeq 2500 and 4000 platforms.

Cutadapt v2.9 (https://doi.org/10.14806/ej.17.1.200) was used to remove sequencing adapters and filter low quality reads (-q 10). Quantification of sequencing reads was performed using Salmon v1.2. ^9^ to the hg38 genome with GC-bias and sequence-specific parameters enabled (--gcbias and –seqbias, respectively), and tximport v1.12.3 (10.12688/f1000research.7563.1) was used to summarize transcript-level to gene-level abundance estimates using R v3.6.0.

### Data Availability Statement

The sequencing data underlying this article are available in public sequencing data from GEO, sratoolkit v2.10.1 (http://ncbi.github.io/sra-tools/). The remaining data underlying this article are available in the article and in its online supplementary material.

### RNA analysis

Raw sequencing counts from Salmon were DESeq2 normalized and VST transformed^10^. Box plots were generated using ggplot2, and PCA was performed using the prcomp function in R v3.6.0.

### Immunohistochemistry (IHC)

Human ileum and colon tissue biopsies from NIBD, colon-like Crohn’s disease (CL) and ileum-like Crohn’s disease (IL) were fixed in 10% (vol/vol) neutral buffered formalin, embedded in paraffin, and prepared as histological sections. After deparaffinization and epitope retrieval in 1X citrate buffer solution, sections were blocked for 1 h in 3% BSA before immunostaining was performed. Polyclonal goat anti-ACE2 antibody (R&D Systems #AF933) was applied overnight at 4^0^C, followed by a 1 h incubation with a secondary anti-goat (Alexa Fluor 594) antibody the next day. Slides were then incubated with DAPI (Invitrogen #D1306) for 5 min to stain nuclei and mounted using FluorSave Reagent (EMD Millipore #345789). Fluorescence was detected using an Olympus VS120 virtual slide microscope.

### ACE2 Signal Intensity

ACE2 fluorescent signal intensity was measured using ImageJ software and normalized to background. To facilitate measurements, images of the stained tissue sections were converted to black and white images on the ACE2 channel, removing signal from DAPI. For each section, pixel intensity was measured in three different regions that were selected for optimal histological cut, showing intact villi (ileum) or colonocytes (colon). Five intensity measurements (e.g., yellow rectangles on supplemental data images) were analyzed per region (**Supplemental Figure 1**). N=4 patients per group. Intensity measurements were averaged per patient and normalized to disease-control group. Significance was determined via one-way ANOVA with multiple comparisons.

### Reverse-Transcriptase qPCR Analysis

Total RNA was extracted from dissected colonic mucosa stored in RNAlater using TRIzol reagent and purified with the Total RNA Purification Plus Kit (48300; Norgen Biotek) according to the manufacturer’s instructions. Total RNA was extracted from isolated and cultured colonic IECs using the Single Cell RNA Purification Kit (51800; Norgen Biotek). Complementary DNA for mRNA was generated from 500 ng of RNA using the High-Capacity Complementary DNA Reverse Transcription Kit (4368814; Thermo Fisher Science). Comparative-Ct-TaqMan with a relative quantification qPCR for mRNAs was performed on the QuantStudio 3 RT-PCR system using TaqMan Fast Advanced Master Mix (4444557; Thermo Fisher Science) with individual TaqMan probes (TaqMan Gene Expression assays, assay ID: Hs01085333_m1 [ACE2], Hs02339424_g1 [RPS9]). Expression of *ACE2* was normalized to *RPS9*.

### Statistical Analysis

All numeric data in the figures are expressed as means ± standard deviation (SD). Differences between the 2 groups were analyzed by a Mann–Whitney or Fisher exact test. Differences between the 3 groups were analyzed by a Kruskal-Wallis test followed by Dunn’s multiple comparison test (R v4.0.1). *P* values less than .05 were considered significant. The Kaplan– Meier method was used to generate survival curves and differences between 2 groups were evaluated by a log-rank test. GraphPad Prism (v8.0; GraphPad Software) was used for these data analyses. Calculation of propensity score and a Cox regression analysis were performed using R v3.5.2.

### Ethical Statement

This study was conducted in accordance with the Declaration of Helsinki and Good Clinical Practice. The study protocol was approved by the Institutional Review Board at the University of North Carolina at Chapel Hill (approval numbers: 19-0819 and 17-0236). All participants provided written informed consent before inclusion in the study. All participants were identified by number and not by name or any protected health information.

All authors had access to the study data and reviewed and approved the final manuscript.

## Results

### ACE2 stratifies two different molecular subtypes of Crohn’s disease

The clinical presentation and course of CD is highly variable. Previously, we found that gene expression data from <u>non-inflamed colon tissue</u> from adult CD (N=28) and non-IBD (NIBD) patients (N=14) clearly segregate CD patients into two disease subtypes ^8^. CD patients in one class largely maintained gene expression profiles of the normal colon (colon-like; CL), whereas in colons of patients in the other class, several normally ileum-specific genes showed robust expression (ileum-like; IL). Altered chromatin accessibility ^8^ and microRNA expression ^11^ across these classes indicated substantive gene regulatory changes, reflecting a fundamental shift in underlying molecular phenotypes. Interestingly, *ACE2, TMPRSS2*, and *TMPRSS4* expression were not significantly different between adult CD and non-IBD (NIBD) patients when considering all CD patients. However, *ACE2* was elevated and *TMPRSS2* and *TMPRSS4* were decreased significantly in IL CD patients relative to CL CD patients (**Figure. 1**). In particular, RNA-seq data showed *ACE2* mRNA levels were 22-fold <u>higher</u> in IL vs CL. Therefore, for the purpose of this paper, we will refer to the two molecular subtypes here as ACE2-high (IL) and ACE2-low (CL).

**Figure 1.**
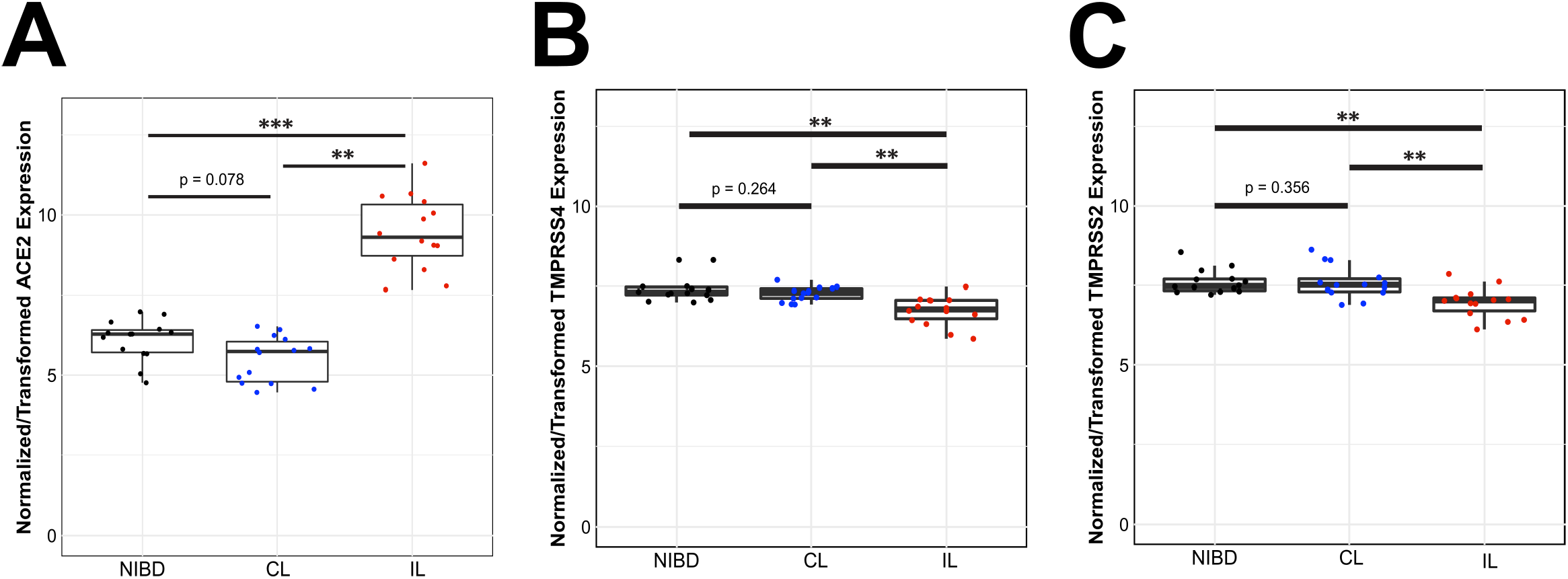
Molecular subtypes of colonic CD. Ileum-like CD (IL) patients express ACE2 and other key marker genes at significantly higher levels than in colon-like CD (CL) patients (*Adj. P< 0.05; **Adj. P<0.005;*** Adj. P<1×10^−6^).

Steady state mRNA expression does not necessarily correspond to protein levels, and transcriptomic assays in bulk tissue lack information on tissue localization. To define the expression and localization of ACE2 in CD at high resolution in intact tissue samples, we performed immunohistochemistry (IHC) on matched formalin-fixed, paraffin-embedded (FFPE) uninflamed colon and ileum tissue from 8 CD and 4 NIBD patients. When divided into CD subclasses based on colonic *ACE2* mRNA expression, ACE2-high CD patients exhibited significantly more ACE2 protein signal compared to NIBD and ACE2-low CD patients (**Figure 2**). Furthermore, abundant immunoreactivity was displayed in villus enterocytes of ileal tissue with a noted significant difference in ACE2 protein between NIBD and ACE2-high CD patients (**Figure 2**). Therefore, we can establish that ACE2 protein levels are strongly correlated with *ACE2* mRNA expression in non-inflamed tissue in our patient cohort, in both the colon and ileum.

**Figure 2.**
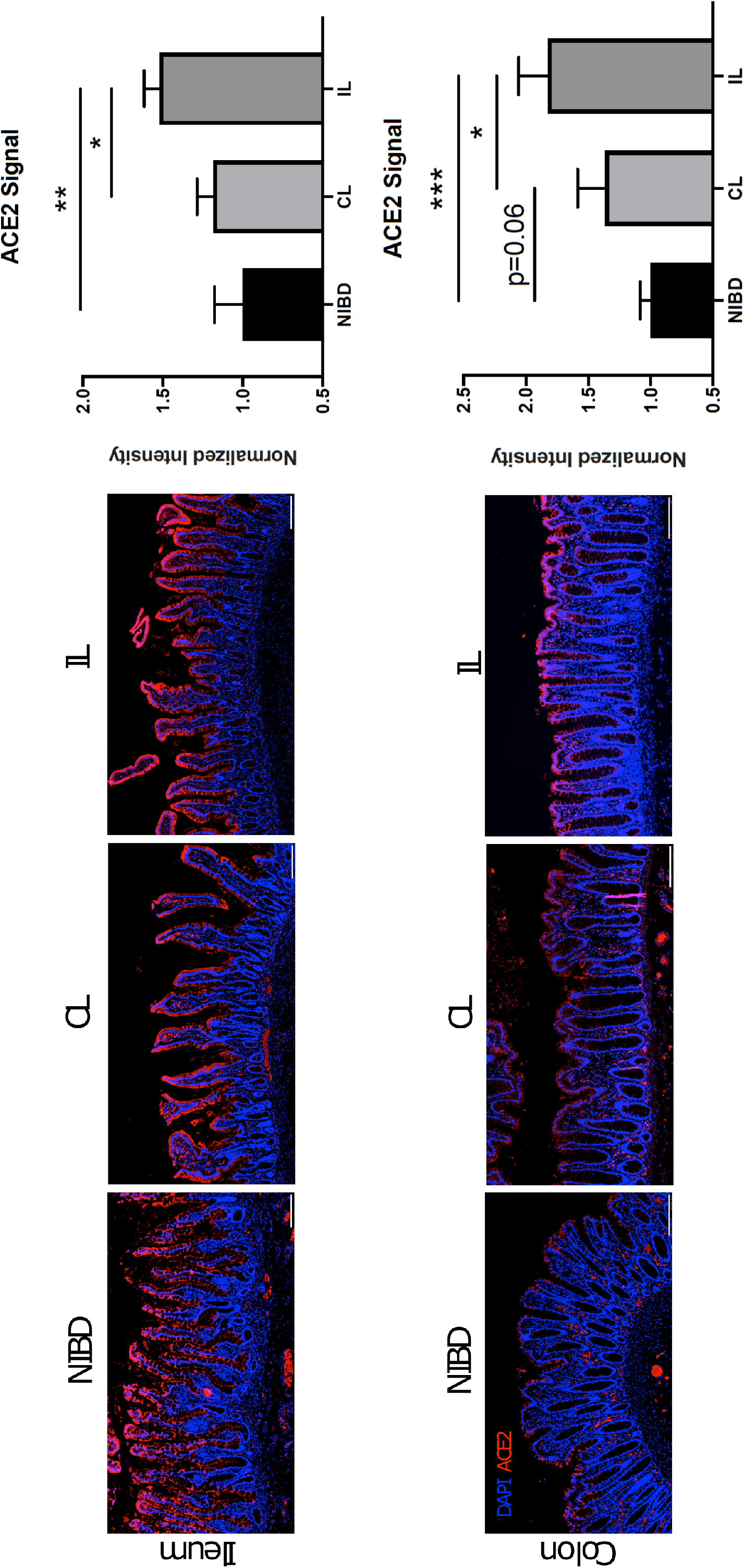
ACE2 IHC reveals increased expression in IL vs CL CD patients. Matched human ileum and colon tissue biopsies from non-IBD (NIBD), CL, and IL patients were stained for anti-ACE2 antibody (pink). Slides were then incubated with DAPI (blue), and ACE2 fluorescent signal intensity was measured using the ImageJ software and normalized to background. N=4 patients per group. Intensity measurements were averaged per patient and normalized to NIBD group. Significance was determined via one-way ANOVA with multiple comparisons. *Adj. P<0.05, **Adj. P<0.005, ***Adj. P<0.001.

### RNA-seq analysis reveals ACE2-high and ACE2-low subclasses in treatment-naïve pediatric Crohn’s disease patients

*ACE2* expression profiles in adult CD patients may vary due to patient treatment histories. Therefore, we sought to determine whether treatment-naïve pediatric CD patients also segregated into similar molecular classes. We performed a principal component analysis (PCA) using our expression data from adult colon samples combined with previously published pediatric expression data from ileal biopsies in age-matched pediatric CD (n=201) and NIBD (n=40) patients generated within the Pediatric Risk Stratification Study (RISK) ^12^ (**Figure 3A**) and analyzed the RISK samples for ACE2 expression levels. Unsurprisingly, samples predominantly separated by study (first principal component). However, two molecular subclasses were evident along the second principal component, similar to the first principal component in single cohort PCAs. Further, *ACE2* expression was highly correlated with the second principal component in the pediatric CD samples (**Figure 3B**), aligning well with the ACE2-high and ACE2-low subclasses defined by our adult CD colon expression data.

**Figure 3.**
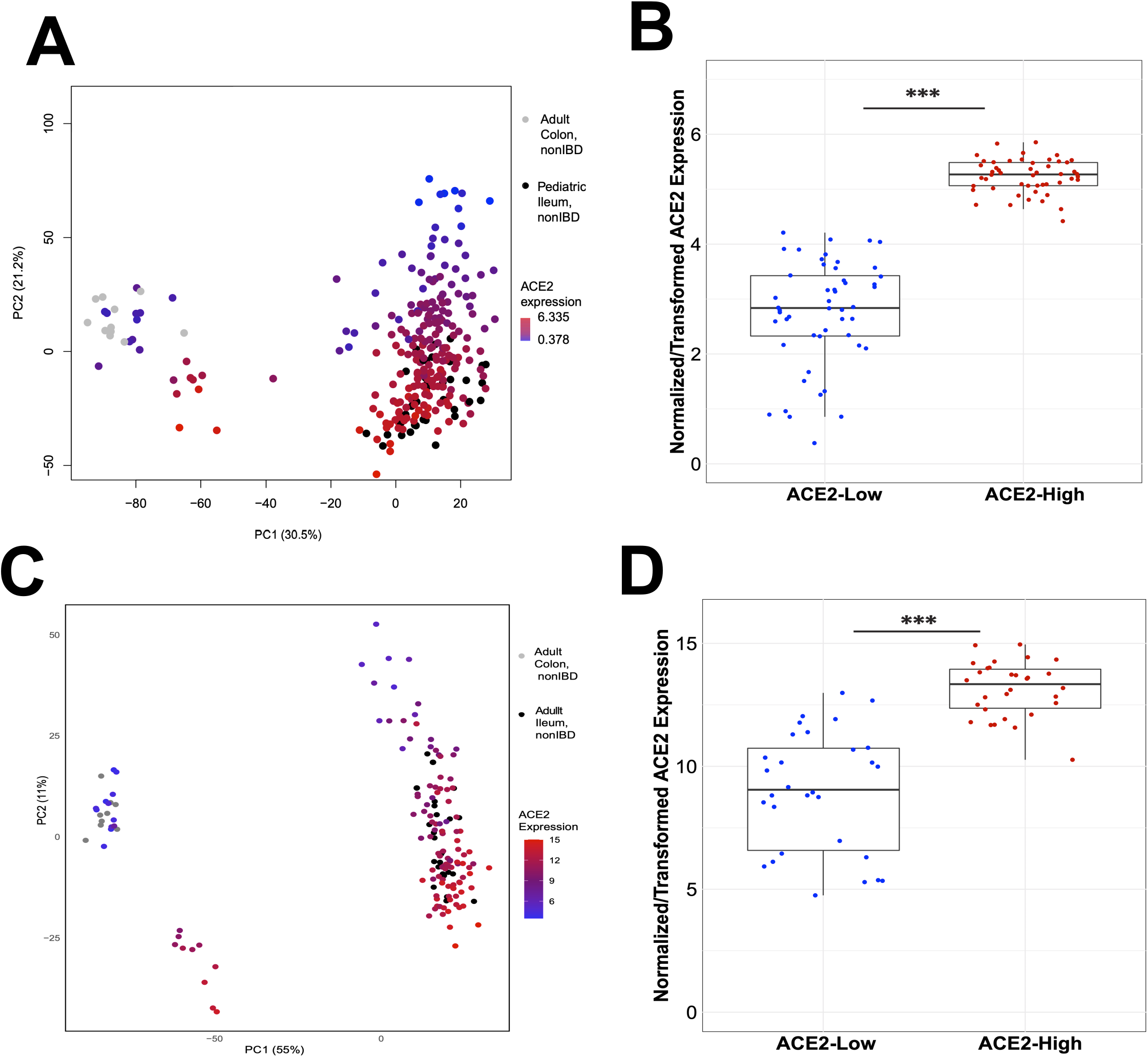
Independent cohorts of adult CD and treatment-naïve pediatric CD ileum samples show similar molecular subtypes. (A) PCA of combined RNA-seq data from adult colon tissue and pediatric ileum tissue from CD and NIBD patients replicates ACE2-High and ACE2-low (PC2) subtypes. (B) Patients from the two extremes of PC2 (panel A; N=50, each direction) show significantly different Ileal expression levels of ACE2 (P<1×10^−6^). (C) PCA of combined RNA-seq data from another independent cohort of adult colon tissue and adult ileum tissue from CD and NIBD patients also replicates ACE2-High and ACE2-low (PC2) subtypes. (D) Patients from the two extremes of PC2 (panel C; N=30 each direction) show significantly different Ileal expression levels of ACE2 (P<1×10^−6^).

Additionally, we performed a combined PCA using our adult samples with expression data from a second previously published study of adult and pediatric ileal biopsies from NIBD (n = 25, no intestinal inflammation and normal histology) and CD (n = 93) patients ^13^ (**Figure 3C**). Again, we observed evidence of ACE2-high and ACE2-low subtypes of CD in this independent cohort of patients (**Figure 3D**).

### Adult colonic and pediatric ileal ACE2 levels correlate with poor clinical outcomes in Crohn’s disease patients

To determine the clinical impact of colonic *ACE2* expression in CD patients, we compared outcomes between 14 ACE2-high and 14 ACE2-low CD patients from the original RNA-seq dataset (**Figure 1**). At the time of CD diagnosis, the only significant difference in clinical characteristics between the subgroups was a higher proportion with ileal involvement in ACE2-high CD patients (92.9 vs. 50.0%, p=0.03; **Supplementary Table 1**). We note that the time to first surgery (bowel resection) within 5 years after CD diagnosis was substantially (but not quite significantly) higher in ACE2-high CD patients (78.6 vs. 42.9%, p=0.12). To better understand this potential relationship to surgery, we generated a Kaplan-Meier plot and performed a subsequent log-rank analysis that showed there was a near-significant difference in time to surgery between the CD subgroups (p=0.095; **Supplementary Figure 2**). To further elucidate the impact of colonic *ACE2* expression on time to first surgery after CD diagnosis, we next performed Cox regression analysis to account for other variables. Covariates shown in **Table 1** and use of anti-TNF alpha agents within 5 years after CD diagnosis were balanced by propensity score for this analysis^11^. Again, we observed a higher risk of surgery in ACE2-high CD patients (OR 3.11; 95%CI, 0.92-10.50; p=0.067; **Supplementary Table 2**).

**Table 1.** Clinical characteristics at the time of CD diagnosis. P values were determined by Mann-Whitney test^a^ or Fisher’s exact test^b^.

|  | ACE2-low | ACE2-high | P value |
| --- | --- | --- | --- |
| Number of patients | 49 | 18 |  |
| Age in years, mean (SD) | 24.0 (11.1) | 27.7 (12.8) | 0.3260<sup>a</sup> |
| Gender, Female (%) | 35 (71.4) | 11 (61.1) | 0.5733<sup>b</sup> |
| Ileal involvement (%) | 37 (75.5) | 17 (94.4) | 0.0936<sup>b</sup> |
| L4 disease (%) | 7 (14.3) | 2 (11.1) | >0.9999<sup>b</sup> |
| Perianal disease (%) | 15 (30.6) | 2 (11.1) | 0.0425<sup>b</sup> |
| Presence of fistula or penetration (%) | 41 (83.7) | 15 (83.3) | >0.9999<sup>b</sup> |
| Current smoking (%) | 14 (28.6) | 8 (44.4) | 0.2504<sup>b</sup> |

To account for a potential Type 1 error due to the small number of samples in the Cox analysis, we analyzed an independent cohort of adult CD patients. Colonic *ACE2* expression was determined in 39 adult CD patients by qPCR. We also performed qPCR on 15 of the 28 CD samples and the 8 of the 14 NIBD from the original cohort (**Figure 1**) to help stratify samples based on ACE2 levels using qPCR. From this unknown cohort, we determined 4 additional patients to be ACE2-high CD patients and 35 ACE2-low CD patients (**Figure 4A**). After combining with our original dataset for a total of 49 ACE2-low and 18 ACE2-high CD patients, we found that the difference in ileal involvement between the two CD subgroups was no longer significant (**Table 1**). However, notably, we did find that ACE2-high CD patients show a significantly higher rate of surgery within 5 years after CD diagnosis than ACE2-low patients (77.8 vs. 44.9 %, p=0.03). Additionally, a Kaplan-Meier analysis revealed a significant difference in the time to first surgery between the two CD subgroups (p=0.04; **Figure 4B**), and a Cox regression analysis found that being in the ACE2-high CD subclass was a significant independent risk factor for surgery (OR 2.27; 95%CI, 1.07-4.80; p=0.032; **Table 2**). Taken together, in this study we discovered that ACE2 expression (mRNA and protein) stratifies two distinct molecular subtypes of CD and that the patients in the ACE2-high subtype have a significantly greater risk of worse clinical outcome and eventual surgery.

**Figure 4.**
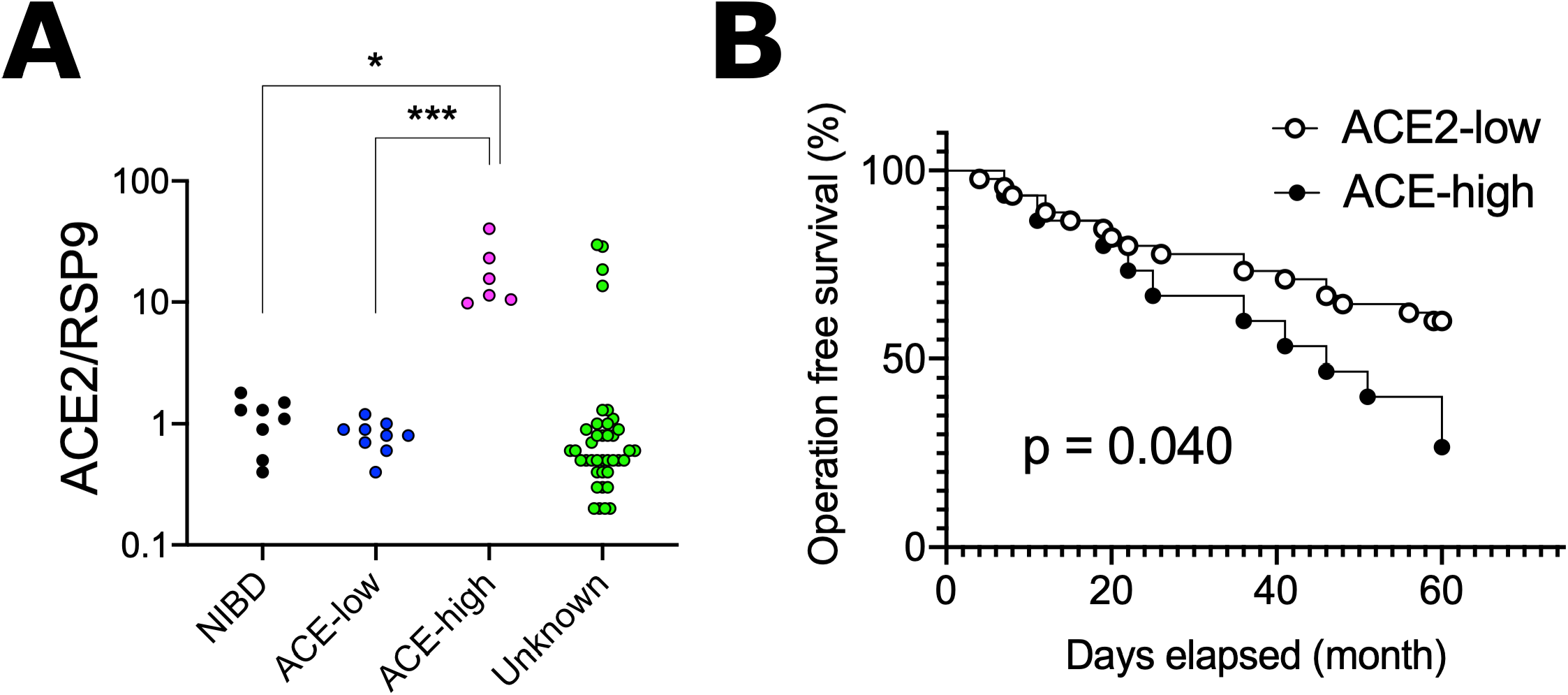
Increased colonic *ACE2* expression is associated with a higher risk of surgery in CD patients. **(A)***ACE2* expression was quantified in colonic specimens obtained from 39 additional CD patients (‘Unknown’ ACE2 expression levels) by qPCR and compared with those from 8 NIBD controls, 8 ACE-low CD, and 6 ACE-high CD patients. **(B)** Kaplan-Meier survival analysis for the risk of surgery within 5 years after CD diagnosis in combined 49 ACE-low and 18 ACE-high CD patients. *p<0.05, ***p<0.001. P-values were determined by Kruskal-Wallis test followed by Dunn’s multiple comparison test (A) and a log-rank test (B).

**Table 2.** Cox logistic regression analysis for surgery. Standard Deviation, SD; odds ratio, OR; confidence interval, CI.

| Variables | Regression<br>Coefficient | SD | P value | OR | 95% CI |
| --- | --- | --- | --- | --- | --- |
| ACE2-high subclass | 0.82 | 0.38 | 0.032 | 2.27 | 1.07 to 4.80 |
| Propensity Score | -0.34 | 1.24 | 0.780 | 0.78 | 0.06 to 8.10 |

## Discussion

Angiotensin-converting enzyme 2 (ACE2) has been thrust into the limelight given its role as a receptor for SARS-CoV-2, responsible for the current COVID-19 pandemic. ACE2 is the key effector peptide of the renin-angiotensin system, mediates vasoconstriction and sodium and water retention both directly and indirectly by stimulating aldosterone secretion. While the impact of ACE2 activity on response to infection is still under debate because no direct evidence has been reported, it is implicated in the response to inflammation and regulation of tissue repair in many organs. ^14, 15^

A recent single cell (sc) RNA-seq study demonstrated that the ACE2-positive-cell ratio along the intestinal tract was significantly higher than in the lung ^14^. In the lung, co-morbidities dramatically increase alveolar *ACE2* expression and are associated with poor outcomes^16^. Furthermore, disease location is a critical determinant of intestinal expression of ACE2 (proteinatlas.org). We reveal through generation and analysis of adult colon RNA-seq data and joint analysis with published adult and pediatric ileal RNA-seq data sets in CD patients that expression of *ACE2* defines two molecular phenotypes of CD, the ACE2-low and ACE2-high patient subsets. Using IHC in matched colon and ileum samples, we validate ACE2 protein expression in apical colonocytes and villus enterocytes for these two patient subsets, as well as NIBD patients.

Our longitudinal analysis from time of CD diagnosis revealed that ACE2-high adult CD patients were associated with increased risk for surgery in the first 5 years after diagnosis. Interestingly, in contrast to worse outcomes in ACE2-high colon expressing adults, reduced ileal *ACE2* expression in CD patients (N=50) in the Pediatric RISK Stratification cohort was significantly associated with colon and ileum disease involvement (p=0.0014), deeper ileal ulcers (p=0.0002), and macroscopic ileal inflammation (p=0.0156) compared to pediatric ileal ACE2-high (N=50) patients. These regional differences in ACE2 expression suggest that active intestinal inflammation alters ACE2 expression, with opposing regulation in ileum and colon^17, 18^. There are several points that must be noted with regard to future comparison of our findings with other studies. First, patient selection is critical as is tissue of origin for analysis, including the inflammatory state of the tissue, which can impact gene expression and interpretation of the results ^5^. Second, *ACE2* expression in the intestine increases with age, making it important to critically evaluate its role separately in different age ^5^groups. Finally, while differences between IBD and NIBD is important, our molecular stratification of CD patients allows for the investigation of two distinct molecular subtypes linked to different clinical phenotypes.

In the small intestine, inflammation and the specific anatomical location were also shown to influence expression of *ACE2* in patients with IBD^4^. We and others showed that in all intestinal segments, *ACE2* expression is much higher in intestinal epithelial cells (IECs) compared to other cells types^19^ (proteinatlas.org). Therefore, understanding how variation in *ACE2* expression and function in IECs impacts IBD activity and COVID-19 severity in IBD patients is critical for managing these individuals. Data from the ongoing SECURE-IBD registry (covidibd.org) indicates unsurprisingly that corticosteroid use increases the risk of severe COVID-19 outcomes >5-fold in IBD patients. Recently, Lukin et al. showed within an inpatient IBD cohort that severe sequelae of COVID-19 were lower than in matched non-IBD controls suggesting a protective effect ^20^. It remains to be seen if variable colonic or ileal ACE2 expression in response to IBD therapeutics is responsible for these observations.

The biological mechanisms impacted by ACE2 in the intestine remain largely unknown and warrant further study. ACE2 functions in the renin-angiotensin system (RAS), counterbalancing the deleterious effects of angiotensin II on the cardiovascular system^21^. Intestinal ACE2 is a chaperone for the amino acid transporter B^0^AT1, a complex in IECs which regulates the gut microbiota^21^. Gut microbiota composition and function, particularly the presence and activity of bacterial and viral pathogens, greatly influence local and systemic immune responses in IBD^22^. Mechanisms driving expression of ACE2 and its co-receptor TMPRSS2 remain unclear. Using an elegant epigenetic approach coupled with genetically manipulated murine models, Chen *et al*., found CDX2, HNF4, SMAD4 and GATA transcription factors bind near *Ace2* and *Tmprss2* resulting in altered chromatin looping and epigenetic modifications with significant impact on *ACE2* and *TMPRSS2* gene expression^19^.

Our present study is significant because we show intestinal ACE2 expression is a biomarker of CD prognosis. Given its well-described link to COVID-19 outcomes in the lung, it is plausible that ACE2 may also serve as a possible injury outcome measure for COVID-19 in IBD. The implications of molecular stratification of CD patients can lead to rapid modification of current therapy in IBD patients impacting the natural course of disease. While actual evidence is still scarce it is hoped that further understanding of the role of ACE2 in IBD pathology and therapeutic responses will ground its use as a biomarker of disease activity and treatment responses contributing to the refinement and development of new therapeutic strategies.

## References

1. Ng SC, Shi HY, Hamidi N, et al. Worldwide incidence and prevalence of inflammatory bowel disease in the 21st century: a systematic review of population-based studies. Lancet 2018;390:2769–2778.

2. Torres J, Mehandru S, Colombel JF, et al. Crohn’s disease. Lancet 2017;389:1741–1755.

3. Ungaro R, Mehandru S, Allen PB, et al. Ulcerative colitis. Lancet 2017;389:1756–1770.

4. Krzysztof NJ, Christoffer LJ, Rahul K, et al. Age, inflammation and disease location are critical determinants of intestinal expression of SARS-CoV-2 receptor ACE2 and TMPRSS2 in inflammatory bowel disease. Gastroenterology 2020.

5. Suarez-Farinas M, Tokuyama M, Wei G, et al. Intestinal inflammation modulates the expression of ACE2 and TMPRSS2 and potentially overlaps with the pathogenesis of SARS-CoV-2 related disease. Gastroenterology 2020.

6. Potdar AA, Dube S, Naito T, et al. Altered intestinal ACE2 levels are associated with inflammation, severe disease and response to anti-cytokine therapy in IBD. Gastroenterology 2020.

7. Furey TS, Sethupathy P, Sheikh SZ. Redefining the IBDs using genome-scale molecular phenotyping. Nat Rev Gastroenterol Hepatol 2019;16:296–311.

8. Weiser M, Simon JM, Kochar B, et al. Molecular classification of Crohn’s disease reveals two clinically relevant subtypes. Gut 2018;67:36–42.

9. Patro R, Duggal G, Love MI, et al. Salmon provides fast and bias-aware quantification of transcript expression. Nat Methods 2017;14:417–419.

10. Love MI, Huber W, Anders S. Moderated estimation of fold change and dispersion for RNA-seq data with DESeq2. Genome Biol 2014;15:550.

11. Keith BP, Barrow JB, Toyonaga T, et al. Colonic epithelial miR-31 associates with the development of Crohn’s phenotypes. JCI Insight 2018;3.

12. Haberman Y, Tickle TL, Dexheimer PJ, et al. Pediatric Crohn disease patients exhibit specific ileal transcriptome and microbiome signature. J Clin Invest 2014;124:3617–33.

13. Mo A, Krishnakumar C, Arafat D, et al. African Ancestry Proportion Influences Ileal Gene Expression in Inflammatory Bowel Disease. Cell Mol Gastroenterol Hepatol 2020;10:203–205.

14. Kurikawa N, Suga M, Kuroda S, et al. An angiotensin II type 1 receptor antagonist, olmesartan medoxomil, improves experimental liver fibrosis by suppression of proliferation and collagen synthesis in activated hepatic stellate cells. Br J Pharmacol 2003;139:1085–94.

15. Donoghue M, Hsieh F, Baronas E, et al. A novel angiotensin-converting enzyme-related carboxypeptidase (ACE2) converts angiotensin I to angiotensin 1-9. Circ Res 2000;87:E1–9.

16. Alqahtani JS, Oyelade T, Aldhahir AM, et al. Prevalence, Severity and Mortality associated with COPD and Smoking in patients with COVID-19: A Rapid Systematic Review and Meta-Analysis. PLoS One 2020;15:e0233147.

17. Burgueno JF, Reich A, Hazime H, et al. Expression of SARS-CoV-2 Entry Molecules ACE2 and TMPRSS2 in the Gut of Patients With IBD. Inflamm Bowel Dis 2020;26:797–808.

18. Verstockt B, Verstockt S, Abdu Rahiman S, et al. Intestinal receptor of SARS-CoV-2 in inflamed IBD tissue seems downregulated by HNF4A in ileum and upregulated by interferon regulating factors in colon. J Crohns Colitis 2020.

19. Chen L, Marishta A, Ellison CE, et al. Identification of Transcription Factors Regulating SARS-CoV-2 Entry Genes in the Intestine. Cell Mol Gastroenterol Hepatol 2020.

20. Lukin DJ, Kumar A, Hajifathalian K, et al. Baseline Disease Activity and Steroid Therapy Stratify Risk of COVID-19 in Patients With Inflammatory Bowel Disease. Gastroenterology 2020;159:1541–1544 e2.

21. Viana SD, Nunes S, Reis F. ACE2 imbalance as a key player for the poor outcomes in COVID-19 patients with age-related comorbidities - Role of gut microbiota dysbiosis. Ageing Res Rev 2020;62:101123.

22. Levy DE, Marie IJ, Durbin JE. Induction and function of type I and III interferon in response to viral infection. Curr Opin Virol 2011;1:476–86.

