## Supplementary figures and images for "Increased Colonic Expression of ACE2 Associates with Poor Prognosis in Crohn’s disease"

### Supplemental Figure 1

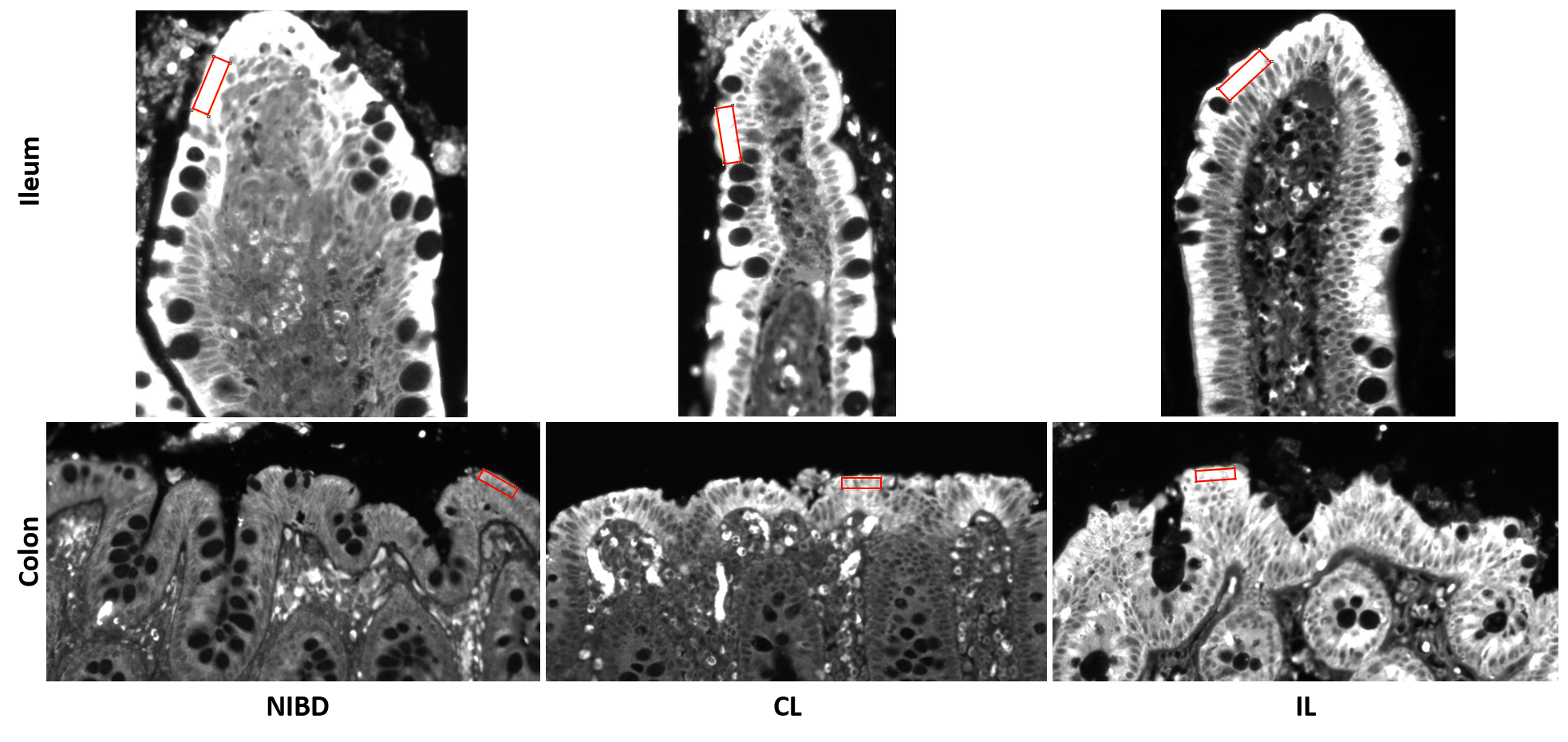
